# Genomic signatures of experimental adaptation to antimicrobial peptides in *Staphylococcus aureus*

**DOI:** 10.1101/023549

**Authors:** Paul R. Johnston, Adam J. Dobson, Jens Rolff

**Affiliations:** Freie Universität Berlin, Königin-Luise-Straße 1-3, 14195 Berlin, Germany; University College London, Darwin Building, Gower Street, London, WC1E 6BT, UK; Berlin-Brandenburg Institute of Advanced Biodiversity Research (BBIB), Altensteinstraße 6, 14195, Berlin, Germany

**Keywords:** Staphylococcus aureus, antimicrobial peptide resistance, experimental evolution

## Abstract

**Objectives:** The evolution of resistance against antimicrobial peptides has long been considered unlikely due to their mechanism of action, yet experimental selection with AMPs results in rapid evolution of resistance in several species of bacteria. Although numerous studies have utilized mutant screens to identify loci that determine AMP susceptibility, there is a dearth of data concerning the genomic changes which accompany experimental evolution of AMP resistance.

**Methods:** Using genome re-sequencing we analysed the mutations which arise during experimental evolution of resistance to the cationic AMPs iseganan, melittin and pexiganan, as well as to a combination of melittin and pexiganan, or to the aminoglycoside antibiotic streptomycin.

**Results:** Analysis of 17 independently replicated *Staphylococcus aureus* selection lines, including unselected controls, showed that each AMP selected for mutations at distinct loci. We identify mutations in genes involved in the synthesis and maintenance of the cell envelope. This includes genes previously identified from mutant screens for AMP resistance, and genes involved in the response to AMPs and cell-wall-active antibiotics. Furthermore, transposon insertion mutants were used to verify that a number of the identified genes are directly involved in determining AMP susceptibility.

**Conclusions:** Strains selected for AMP resistance under controlled experimental evolution displayed consistent AMP-specific mutations in genes which determine AMP susceptibility. This suggests that different routes to evolve resistance are favored within a controlled genetic background.

## Introduction

Antimicrobial peptides (AMPs), ubiquitous in multicellular organisms^1^, are considered to be a promising source of new and potent antibiotics^2^. Current research on AMPs mostly focuses on the mechanisms of action and on the development of therapeutics whereas only a small number of studies have addressed the important problem of bacterial resistance evolution. Resistance against cationic AMPs evolves readily *in vitro* in *Escherichia coli* and *Pseudomonas aeruginosa*^3^, *Salmonella enterica*^4^, and *Staphylococcus aureus^5,6^.* Experimentally evolved strains of *S. aureus* that were selected successfully for resistance against the catioinc protegrin-1 analog iseganan^6^ survive better in a model host^7^, which relies heavily on AMPs to deal with long-lasting infections^8^. *S. aureus* populations selected for resistance to pexiganan and mellitin also show a trend towards increased survival in the host^7^. Here we present a genomic analysis of *S. aureus* strains from these populations^6^ together with susceptibility data from transposon insertion mutants that show a number of the identified genes are directly involved in mediating AMP susceptibility.

## Materials and methods

Strains were isolated from populations which were created by selecting *S. aureus* JLA513^9^ (*hla-lacZ hla*+, derived from SH1000, from Simon Foster, University of Sheffield) for 28 days with increasing concentrations of AMPs or with the aminoglycoside antibiotic streptomycin^6^. Streptomycin-selected strains are included here as a positive control since the genetic basis of streptomycin resistance is well-characterized in *S. aureus*. Briefly, to ensure adaptation to the culture medium 50 μl of *S. aureus* JLA513 culture was passaged serially every 24 h for 10 days in 5 ml Müller-Hinton Broth (MHB). Subsequently, 5 parallel selection lines were established in each treatment at MIC_50_ (as well as unselected controls) by innoculating 5 μl of serially-passaged culture into 500 μl of MHB containing the cognate selective agent. 5 μl of 24 h cultures were passaged daily to fresh MHB. The concentrations of the selective agents were doubled each week for a total of four weeks. See Dobson et al. 2013 Table S1 for full details and precise concentrations^6^. Strains were isolated from each of three independently selected replicate populations per selective agent (with the exception of iseganan-selected populations where only 2 frozen population stocks remained viable), as well as from unselected controls and the ancestral strain JLA513. Minimum inhibitory concentrations (MIC) were calculated for the selective agents (Table S1) in 96-well plates as previously described^10^ and DNA was isolated from each strain using a Roboklon DNA extraction kit (Roboklon GmbH, Germany). Genomic DNA from each strain was sequenced for 180 cycles using a HiSeq2000 by the Beijing Genomics Institute (BGI), resulting in 90-bp paired-end reads. Sequence data are available from the NCBI SRA under BioProject ID PRJNA291485. Strain JLA513^9^ was constructed using strain SH1000, which is a derivative of strain 8325. The genetic differences between SH1000 and other members of the 8325 lineage have been described using both array-based resequencing^11^ and subsequently by de novo genome sequencing^12^. The differences comprise: the excision of three prophages from strain 8325 (Φ11, 12, 13), 13 single-nucleotide polymorphisms (2 synonymous, 11 non-synonymous), a 63-bp deletion in the *spa-sarS* intergenic region, and an 11-bp deletion in *rsbU*^*12*^. Therefore a consensus reference genome was first produced to account for these differences. Reads from JLA513 were assembled using SPAdes^13^ and the resulting contigs were used to correct for the 3 phage excision sites in the 8325 reference genome. JLA513 reads were then mapped to the resulting sequence and bcftools consensus^14^ was used to correct the remaining 13 SNPs and 2 indels. To identify mutations in the selection lines, reads were mapped to this reference genome using BWA^15^ and sorted, deduplicated (to account for optical-and PCR-duplicates) and indexed using SAMtools^14^ and Picard (http://broadinstitute.github.io/picard). Average coverage was 134-fold (range 110-144 fold). Variants were called using FreeBayes version v0.9.14-8-g1618f7e^16^ and coverage was calculated across 25-bp windows using IGVtools^17^. All variants were independently verified using a second computational pipeline, breseq^18^. Insertion mutants were obtained from the Nebraska Transposon Mutant Library^19^ in order to test if the identified genes were directly involved in AMP resistance. MICs were calculated for each mutant and the wild type strain USA300_FPR3757 as described above.

## Results and discussion

Between one and four mutations were identified per strain after accounting for differences between the JLA513 ancestor and the 8325 reference genome, and for mutations arising over the course of the experiment across treatments and unselected controls. In total, 28 mutations were identified across the 17 strains including 24 nonsynonymous mutations affecting 13 genes, a segmental duplication of 124-kb region containing an entire *rrn* operon (Table 1, Table S2) as well as 1 synonymous mutation and 2 intergenic indels (Table S2).

Pexiganan resistance was characterized by distinct nonsense mutations in the gene encoding the XRE-family transcriptional regulator XdrA in strains PG2.2 and PG4.2 (Table 1, Table S2). XdrA was recently shown to activate transcription of *spa*^20^, which encodes the protein A virulence factor, and deletion mutants show increased β-lactam resistance^21^. Here, a transposon mutant with an insertion in *xdrA* showed decreased pexiganan susceptibility (Table 1, Table S3) indicating that XdrA is directly involved in pexiganan resistance. In addition to a mutation in *xdrA*, strain PG4.2 also carried a nonsynonymous substitution in *wcaG*, which encodes a putative UDP-glucose-4 epimerase (Table 1). Only a single mutation was observed in strain PG1.1, introducing a frameshift into *mgt* (*sgtB*), which encodes a monofunctional peptidoglycan glycosyltransferase (Table 1). A distinct nonsense in *mgt* was also identified in one pexiganan-melittin-selected strain (see below). An *mgt* transposon mutant was also found to be less susceptible to pexiganan (Table 1, Table S3). As part of the cell wall stimulon^22^, *mgt* is positively regulated by cell wall stress and participates in the polymerization of lipid II into nascent peptidoglycan^23^. Recent work has shown that *mgt* mutations cause peptidoglycan chain length reduction as well as alterations in cellular morphology and division site placement^24^.

**Table 1.** Mutations identified in strains selected for resistance to different antimicrobials.

| Selection | No. of strains<sup>a</sup> | Gene | Function | Locus tag<sup>b</sup> | Susceptibility of Tn mutant<sup>c</sup> |
| --- | --- | --- | --- | --- | --- |
| IG | 2 | *yjbH* | Disulfide stress response | SAOUHSC_00938 | not tested |
| ML | 1 | *walR (yycG)* | Cell envelope biogenesis | SAOUHSC_00020 | not tested |
| ML | 3 | *rluA* | Pseudouridine synthase | SAOUHSC_00944 | unchanged |
| ML/PGML | 3(2ML/1PGML) | *ytrA* ortholog | Cell wall stimulon | SAOUHSC_02155 | not tested |
| PG | 1 | *wcaG* | Nucleoside-diphosphate-sugar epimerase | SAOUHSC_00664 | unchanged |
| PG | 2 | *xdrA* | Xenobiotic response element | SAOUHSC_01979 | decreased |
| PG | 2(1PG/1PGML) | *mgt (sgtB)* | Cell wall stimulon | SAOUHSC_02012 | decreased |
| PGML | 1 | *hpr* | Carbohydrate transport | SAOUHSC_01028 | not tested |
| PGML | 1 | *dak2* | Cell envelope biogenesis | SAOUHSC_01193 | increased |
| PGML | 1 | *putA (fadM)* | Amino acid metabolism | SAOUHSC_01884 | unchanged |
| STR | 1 | *nusA* | Transcription antitermination | SAOUHSC_01243 | not tested |
| STR | 2 | *glpK* | Glycerol kinase | SAOUHSC_01276 | unchanged |
| STR | 1 | *rrn* operons | Ribosome biogenesis | 124-kb *rrn* region | not tested |
| STR | 3 | *gidB (rsmG)* | Ribosome biogenesis | SAOUHSC_03051 | decreased |

<sup>a</sup> Number of strains with a mutation in a given gene.
<sup>b</sup> Identifier in *Staphylococcus aureus* NCTC 8325 reference genome.
<sup>c</sup> Susceptibility of transposon insertion mutants from the Nebraska Transposon Mutant Library to the cognate selective agent. Not tested, transposon mutant not available. See Table S3 for full details.
IG, iseganan; ML, melittin; PG, pexiganan; PGML, 1:1 wt/wt combination of melittin and pexiganan; STR, streptomycin. See Table S4 for further details on AMPs used.

All 3 melittin-resistant strains were found to carry missense mutations resulting in either A35T or A35D substitutions in a gene encoding a putative RluD-like pseudouridylate synthase with no known role in antimicrobial susceptiblity. A transposon mutant from the Nebraska Transposon Mutant Library with an insertion in this gene showed no change in melittin susceptibility (Table 1, Table S3). One melittin-resistant strain carried a L93I missense mutation in a region encoding an alpha helix immediately adjacent to the conserved active site quintet in the response regulator WalR (Table 1). WalKR regulates cell wall metabolism and is ubiquitous in the *Firmicutes* where it is the only known essential two-component system^25^. *walKR* mutations, including those affecting the WalR active site, arise during persistent clinical *S. aureus* infections and are known to confer resistance to vancomycin and the lipopeptide antibiotic daptomycin by increasing the thickness of the cell wall^26^. Identical nonsense mutations were idenitified in two melittin-resistant strains at the extreme 5′ end of the *ytrA* open reading frame, which encodes a winged helix-turn-helix GntR-family repressor (Table 1). Similar to its *B. subtilis* ortholog, *ytrA* is the first gene of an operon which encodes two putative ABC transporters. In *B. subtilis*, YtrA binds specifically to an inverted repeat in the *ytrA* and *ywoB* promoters, and transcription of the *ytr* and *ywo* operons is induced by cell-wall-active antibiotics including the peptide antibiotics bacitracin, vancomycin and ramnoplanin, with *ytrA* null mutations causing constitutive expression of both operons^27^. Notably, the entire *ytrA* operon has been shown to be induced by cationic AMPs in *S. aureus*, where it is under negative regulation by the AMP sensing system *aps*^*28*^ and has also been implicated in nisin susceptibility in *S. aureus* SH1000^29^. Although *ytrA* insertions are not present in the Nebraska Transposon Mutant Library we were able to obtain 2 independent *ytr* operon transposon mutants with insertions downstream of *ytrA* which did not show any detectable difference in AMP susceptiblity relative to the wild type (Table S3). This raises the possibility that the *ytrA*-null mutations observed here may mediate AMP susceptibility via derepression of the *S. aureus ywo* ortholog.

Iseganan resistance was associated with an identical 5-bp deletion in the extreme 3′ end of the *yjbH* gene in each of two strains from independent iseganan-selected lines (Table 1). YjbH controls the disulfide stress response by binding to the oxidative burst-specific transcriptional regulator Spx, and thereby controlling its degradation by the ClpXP protease^30^, a role which is conserved in *Bacillus subtilis*^31^. YjbH also modulates ß-lactam susceptibility, with deletion mutants showing moderate resistance to various ß-lactams but not to the glycopeptide antibiotic vancomycin^30^. The precise mechanism by which YjbH modulates susceptibility is unknown but is proposed to be a consequence of upregulation of PBP4 which results in increased peptidoglycan cross-linking^30^.

There were no common mutations identified in the genomes of three strains which were selected with a 1:1 wt/wt combination of pexiganan and melittin (Table 1). However there were commonalities with strains that were selected with either melittin or pexiganan. A single missense mutation was identified in strain PGML3.2 which substitutes a conserved threonine residue in the winged helix-turn-helix DNA binding domain of YtrA (note that *ytrA* nonsense mutations were identified in 2 melittin-resistant strains described above). Similarly, a single nonsense mutation was identified in strain PGML5.1 in *mgt* (also mutated in 1 pexiganan-resistant strain described above). In contrast, three missense mutations were identified in the genome of a second pexiganan-melittin-selected strain. Interestingly this included *dak2* which encodes a dihydroxyacetone kinase responsible for incorporation of diphosphatidylglycerol into the cell membrane^32^. *dak2* was previously identified in a high throughput mutant screen for loci affecting susceptibility to the anionic human AMP dermcidin in *S. aureus^32^*. Mutations affecting the non-essential C-terminal DegV superfamily domain of Dak2 result in altered membrane phospholipid composition and decreased binding and activity of dermcidin but not of the catioinic human AMPs LL-37 or human ß-defensin-3^32^. Given this lack of cross-resistance to cationic AMPs in *dak2* mutants, Dak2-mediated susceptibilty was thought to be specific to anionic AMPs such as dermcidin^32^. It is therefore surprising to find *dak2* mutation in response to selection with a combination of the cationic AMPs mellittin and pexiganan. Further evidence of the role of Dak2 in susceptibility to pexiganan and melittin was shown by increased susceptibility to both AMPs by a *dak2* transposon mutant (Table 1, Table S3).

Mutations identified in streptomycin-selected strains mostly occurred in genes with known roles in streptomycin susceptibility (Table 1). Frameshift mutations in *gidB*, which encodes a 16S rRNA-specific 7-methylguanosine methyltransferase, were identified in all three streptomycin-selected strains (Table 1). In each case, the frameshift occurs within the region encoding the GidB methyltransferase domain. Mutations in *gidB* (*rsmG*) are associated with low-level streptomycin resistance in several species of bacteria including *S. aureus^33–36^* and it is speculated that loss of 16S methylation lowers the binding affinity of streptomycin thus conferring the resistance phenotype^35^. Here, a *gidB* transposon mutant was found to be 4-fold less susceptible to steptomycin (Table 1, Table S3). Two further mutations were identified which potentially affect ribosomal RNA. A 124-kb region containing an entire *rrn* operon appears to have been duplicated in a strain STR3.2 whereas strain STR1.1 carries a non-sysnonymous substitution in the essential gene encoding NusA, which acts as an antiterminator for 16S rRNA transcription, as well as a chaperone for 16S rRNA folding^37^ (Table S2). Mutations were also identified in the glycerol kinase gene *glpK* in two strains (Table 1) however a transposon insertion did not detectably alter streptomycin susceptibility (Table 1, Table S3).

Numerous studies have utilized mutant screens to identify loci that determine AMP susceptibility^32,38^ but with the exception of a single study^4^, there is a dearth of data concerning the genomic changes which accompany experimental evolution of AMP resistance. Here, genome sequencing of strains isolated from independently replicated AMP selection lines identified mutations associated with AMP resistance evolution and showed that each AMP selected for mutations at distinct loci. These mutations affected genes with known roles in susceptibility to AMPs and/or cell-wall-active antibiotics, as well as cell wall stress stimulon genes. All cationic AMPs used here form toroidal pores, yet there was little evidence of cross resistance or for mutations that were common across all AMP-selected strains. There is limited evidence of AMP-specific responses. For example, the staphylococcal virulence factor MprF determines susceptibility towards protegrins (e.g. iseganan) but has little effect on magainin (pexiganan analog) or melittin susceptibility^39^. Also, little is known about AMP interactions with other constituents of the cell membrane and whether these may contribute to the specificity observed here. A small number of mutations occurred in genes with no known role in antimicrobial susceptibility, such as the gene encoding the RluD-like pseudouridylate synthase, and may represent compensatory adaptations that warrant further study. Furthermore, mutations in the *walR* gene such as that described here are known to increase multidrug resistance and to arise during clinical *S. aureus* infections^26^. This is consistent with the notion that the evolution of resistance to AMPs may compromise host defences against infection^5,40^.

## Acknowledgements

We are grateful to Christine Bergmann, Sabine Kretschmer, and Uta Müller for technical assistance as well as to Simon Foster, Joshua Hooker, Alex O’Neill, and Christopher Randall for providing materials.

## Transparency declarations

None to declare.

## Supplementary data

Table S1. MICs for various antimicrobials against 18 strains of *S. aureus*.

**TABLE S1.** MICs for various antimicrobials against 18 strains of *S. aureus*.

| Strain | MIC(ug/ml)<sup>a</sup> |  |  |  |  |
| --- | --- | --- | --- | --- | --- |
|  | Melittin | Pexiganan | Pex-Mel<sup>b</sup> | Streptomycin | Vancomycin |
| JLA513 | 8 | 8 | 8 | 4 | 2 |
| IG1.2 | 4 | 8 | 8 | 4 | 2 |
| IG2.1 | 4 | 8 | 8 | 4 | 2 |
| ML1.1 | 32 | 8 | 32 | 8 | 4 |
| ML4.2 | 32 | 8 | 16 | 4 | 2 |
| ML5.2 | 32 | 16 | 16 | 4 | 2 |
| PG1.1 | 8 | 16 | 8 | 4 | 2 |
| PG2.2 | 4 | 16 | 8 | 4 | 2 |
| PG4.2 | 4 | 16 | 8 | 8 | 2 |
| PGML3.2 | 16 | 16 | 16 | 2 | 2 |
| PGML4.4 | 8 | 32 | 16 | 4 | 2 |
| PGML5.1 | 8 | 16 | 16 | 2 | 2 |
| STR1.1 | 8 | 8 | 8 | 32 | 2 |
| STR2.2 | 8 | 8 | 8 | >64 | 2 |
| STR3.2 | 8 | 16 | 8 | >64 | 2 |
| Uns1.1 | 4 | 4 | 4 | 4 | 2 |
| Uns3.4 | 4 | 4 | 4 | 4 | 2 |
| Uns4.2 | 8 | 8 | 8 | 8 | 2 |

<sup>a</sup>MIC, minimum antimicrobial concentration necessary to inhibit the growth of *S. aureus*.
<sup>b</sup>Equal quantities of pexiganan and melittin.
ML, melittin; PG, pexiganan; PGML, 1:1 wt/wt combination of melittin and pexiganan; STR, streptomycin; Uns, unselected control strain.

Table S2. Summary of all mutations.

**TABLE S2.** Summary of all mutations.

| Strain | Mutation | Locus tag<sup>a</sup> | Annotation | Function |
| --- | --- | --- | --- | --- |
| IG1.2 | p.S266IfsX45 | SAOUHSC_00938 | *yjbH* | Disulfide stress response |
| IG2.1 | p.S266IfsX45 | SAOUHSC_00938 | *yjbH* | Disulfide stress response |
| ML1.1 | p.L93I | SAOUHSC_00020 | *walR/yycG* | Cell envelope biogenesis; response regulator |
| ML1.1 | p.A35T | SAOUHSC_00944 | *rluD*-like | Pseudouridylylate synthase |
| ML1.1 | g.2101984_2101985insT | SAOUHSC_02270 | intergenic | - |
| ML4.2 | p.A35D | SAOUHSC_00944 | *rluD*-like | Pseudouridylylate synthase |
| ML4.2 | p.L5X | SAOUHSC_02155 | *ytrA* | Cell wall stimulon; repressor |
| ML5.2 | p.A35D | SAOUHSC_00944 | *rluD*-like | Pseudouridylylate synthase |
| ML5.2 | p.L5X | SAOUHSC_02155 | *ytrA* | Cell wall stimulon; repressor |
| PG1.1 | p.P39XfsX3 | SAOUHSC_02012 | *mgt/sgtB* | Cell wall stimulon; peptidoglycan glycosyltransferase |
| PG2.2 | p.Q40RfsX24 | SAOUHSC_01979 | *xdrA* | Xenobiotic response element |
| PG4.2 | p.M280V | SAOUHSC_00664 | *wcaG* | Nucleoside-diphosphate-sugar epimerase; oxidoreductase |
| PG4.2 | p.Q30X | SAOUHSC_01979 | *xdrA* | Xenobiotic response element |
| PGML3.2 | p.T74A | SAOUHSC_02155 | *ytrA* | Cell wall stimulon; repressor |
| PGML4.4 | p.A16D | SAOUHSC_01028 | *hpr* | Carbohydrate transport |
| PGML4.4 | p.G341D | SAOUHSC_01193 | *dak2* | Cell envelope biogenesis; dihydroxyacetone kinase |
| PGML4.4 | p.S138I | SAOUHSC_01884 | *putA/fadM* | Amino acid metabolism; proline dehydrogenase |
| PGML5.1 | p.Q251X | SAOUHSC_02012 | *mgt/sgtB* | Cell wall stimulon; peptidoglycan glycosyltransferase |
| STR1.1 | p.A227E | SAOUHSC_01243 | *nusA* | Transcription antitermination; antiterminator |
| STR1.1 | p.H87L | SAOUHSC_02727 | NC_007795.1 | Hypothetical protein; peptidase |
| STR1.1 | p.R218DfsX75 | SAOUHSC_03051 | *gidB/rsmG* | Ribosome biogenesis; 16S rRNA methyltransferase |
| STR2.2 | c.63A>G<sup>b</sup> | SAOUHSC_00489 | *folP* | Dihydropteroate synthase |
| STR2.2 | g.1090526_1090533del | intergenic | - | - |
| STR2.2 | p.A332E | SAOUHSC_01276 | *glpK* | Glycerolipid metabolism; glycerol kinase |
| STR2.2 | p.S115EfsX12 | SAOUHSC_03051 | *gidB/rsmG* | Ribosome biogenesis; 16S rRNA methyltransferase |
| STR3.2 | p.G251X | SAOUHSC_01276 | *glpK* | Glycerolipid metabolism; glycerol kinase |
| STR3.2 | g.2122437_2246248dup | segmental duplication | - | Encodes rRNA and ribosomal protein genes |
| STR3.2 | p.S115EfsX12 | SAOUHSC_03051 | *gidB/rsmG* | Ribosome biogenesis; 16S rRNA methyltransferase |

<sup>a</sup>Identifier in *Staphylococcus aureus* NCTC 8325 reference genome.<sup>b</sup>Synonymous.
IG, iseganan; ML, melittin; PG, pexiganan; PGML, 1:1 wt/wt combination of melittin and pexiganan; STR, streptomycin.

Table S3. MICs for various antimicrobials against transposon insertion mutants of *S. aureus* strain USA300_FPR3757 from the Nebraska Transposon Mutant Library.

**TABLE S3.** MICs for various antimicrobials against transposon insertion mutants of *Staphylococcus aureus* strain USA300_FPR3757 from the Nebraska Transposon Mutant Library.

| Strain | Locus tag<sup>b</sup> | Annotation | MIC(ug/ml)<sup>a</sup> |  |  |  |
| --- | --- | --- | --- | --- | --- | --- |
|  |  |  | Melittin | Pexiganan | Pex-Mel<sup>c</sup> | Streptomycin |
| USA300 | - | - | 8 | 16 | 16 | 4 |
| NE229 | SAUSA300_1119 | *dak2* | 8 | **8** | **8** | 4 |
| NE239 | SAUSA300_1711 | *putA* (*fadM*) | 8 | 16 | 16 | 4 |
| NE249 | SAUSA300_2644 | *gidB* (*rsmG*) | 8 | 16 | 16 | **16** |
| NE467 | SAUSA300_0644 | *wcaG* | 8 | 16 | 16 | 4 |
| NE596 | SAUSA300_1855 | *mgt* (*sgtB*) | 8 | **32** | 16 | 4 |
| NE822 | SAUSA300_0909 | *rluD*-like | 8 | 16 | 16 | 4 |
| NE896 | SAUSA300_0903 | *yjbH* | 8 | 16 | 16 | 4 |
| NE1023 | SAUSA300_0984 | *ptsI* | 8 | 16 | 16 | 4 |
| NE1445 | SAUSA300_1797 | *xdrA* | 8 | **32** | **8** | 4 |
| NE1587 | SAUSA300_1192 | *glpK* | 8 | 16 | 16 | 4 |
| NE1908<sup>d</sup> | SAUSA300_1911 | ABC transporter | 8 | 16 | 16 | 4 |
| NE1188<sup>d</sup> | SAUSA300_1912 | ABC transporter | 8 | 16 | 16 | 4 |

<sup>a</sup>MIC(minimum inhibitory concentration), minimum antimicrobial concentration necessary to inhibit the growth of *S. aureus*.
<sup>b</sup>Identifier in *S. aureus* USA300\_FPR3757 reference genome.
<sup>c</sup>Equal quantities of pexiganan and melittin.
<sup>d</sup>Insertions in the *ytr* operon downstream of *ytrA*. Insertions in *ytrA* are not present in the Nebraska Transposon Mutant Library.
ML, melittin; PG, pexiganan; PGML, 1:1 wt/wt combination of melittin and pexiganan; STR, streptomycin.

Table S4. Details of antimicrobial peptides used.

**TABLE S4.**
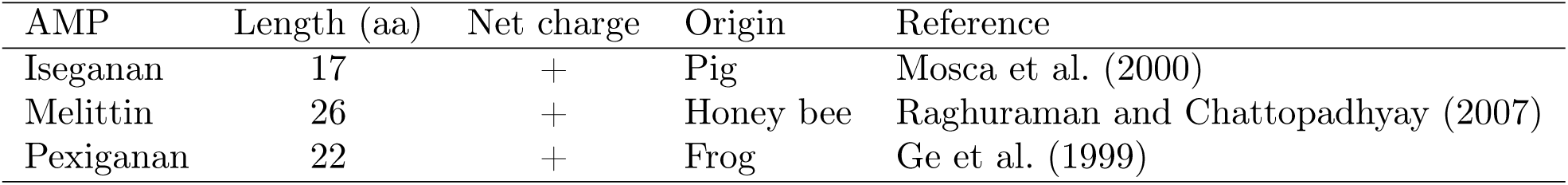
Details of antimicrobial peptides used.

| AMP | Length (aa) | Net charge | Origin | Reference |
| --- | --- | --- | --- | --- |
| Iseganan | 17 | + | Pig | Mosca et al. (2000) |
| Melittin | 26 | + | Honey bee | Raghuraman and Chattopadhyay (2007) |
| Pexiganan | 22 | + | Frog | Ge et al. (1999) |

